# Wayanad Sky Islands Survey 2025

**DOI:** 10.1101/2025.05.29.656751

**Authors:** MP Sivachand, RL Rathish, Ek Shyama, CK Vishnudas

**Affiliations:** Hume Centre for Ecology & Wildlife Biology; Kerala Veterinary and Animal Sciences University

**Author notes:** Research Assistant, Ecosystem & Wildlife, Hume Centre for Ecology & Wildlife Biology (MP Sivachand, Ek Shyama). Principal Investigator, Ecosystem & Wildlife, Hume Centre for Ecology & Wildlife Biology (CK Vishnudas).

**Keywords:** Sky Islands, Shola, Western Ghats, eBird, Wayanad

## Abstract

The Shola Sky Islands of the Western Ghats are isolated montane habitats that harbour significant avian diversity and endemism. This study assessed bird species diversity across five mountain ranges in Wayanad, Kerala (Banasura, Brahmagiri, Camels Hump, Kuricharmala, and Manikunnu) situated above 1200 meters. A total of 114 species were recorded, with the Camels Hump range supporting the highest diversity (81 species). Patterns of species richness were found to correspond with mountain area and degree of isolation, consistent with the principles of island biogeography. Birds were classified by taxonomic order and family, with Passeriformes being the most dominant order. Among families, Muscicapidae recorded the highest number of species, followed by Accipitridae and Phylloscopidae. Invertivores constituted the majority of trophic guilds, and species richness showed a general decline with increasing elevation. These findings highlight the importance of habitat area and connectivity in shaping avian communities within sky island ecosystems. They also underline the urgent need for targeted conservation efforts in this ecologically sensitive region, much of which falls under territorial divisions, including reserve and vested forests.

## 1. INTRODUCTION

The Western Ghats are globally recognised as a crucial biodiversity hotspot. Stretching along India’s western coast, roughly 30 to 50 kilometres inland, this mountain range spans the states of Kerala, Tamil Nadu, Karnataka, Goa, Maharashtra, and Gujarat. Covering approximately 140,000 square kilometres over 1,600 kilometres, the Ghats are interrupted only by the 30-kilometre-wide Palghat Gap and the smaller Shencottah Gap (Myers *et al*.,2000). Among its most remarkable features is the Shola Sky Islands, a unique ecological phenomenon characterised by high elevation, isolation, diverse microclimates, and significant biodiversity. These montane habitats, ranging between 1,200 and 2,400 meters above sea level, are composed of a mosaic of evergreen forests and grasslands, commonly referred to as the shola-grassland complex.

The concept of island biogeography, first articulated by MacArthur and Wilson, posits that species diversity on islands is shaped by island size and distance from the mainland, influencing immigration and extinction rates (Borregaard *et al*., 2016). This theory is particularly relevant to the Shola Sky Islands, where geographical isolation has driven high levels of endemism and unique species assemblages (Gupta *et al*., 2019). The separation of these habitats has fostered distinct evolutionary pathways, leading to the emergence of specialised traits among species, such as unique vocalisations and breeding behaviours adapted to specific habitat conditions (Robin *et al*., 2017; Purushotham & Robin, 2016). However, the ecological integrity of these sky islands faces growing threats from anthropogenic pressures, particularly the introduction of non-native timber species such as Acacia, Eucalyptus, and Pinus (Jobin *et al*., 2023). These invasives outcompete native flora and facilitate further invasions, leading to cascading ecological disruptions (Jobin *et al*., 2022). Over time, the proliferation of plantations has caused significant habitat loss, with tropical montane grasslands reduced by as much as two-thirds in some areas (Arasumani *et al*., 2018). The resulting habitat fragmentation affects not only plant communities but also animal populations, as evidenced by studies on butterfly populations showing reduced gene flow and increased genetic structuring in isolated patches (S & P, 2013). Maintaining habitat connectivity thus becomes vital for species survival and ecosystem resilience in the face of environmental changes.

Located at the highest elevations of the Western Ghats, the Shola Sky Islands are particularly vulnerable to the impacts of global warming, further highlighting the urgency for focused conservation efforts (Robin & Nandini, 2012). In this context, the present study, conducted in regions above 1,200 meters above sea level, aims to examine bird diversity and assess variation across mountain ranges, elevation bands, and habitat types. It also explores trophic niches and dominant bird groups within these habitats. Such research is essential for evaluating the conservation status of these highly biodiverse yet under-protected landscapes, which host endangered species like the Banasura Laughingthrush. Effective conservation strategies are needed to ensure the long-term survival of these exceptional ecosystems.

## 2. METHODOLOGY

The study area focuses on Wayanad’s mountainous and hilly regions, situated on the western edge of the Western Ghats, spanning elevations between 1,200 and 2,100 meters above mean sea level (msl). It extends from the Brahmagiri ranges in the north to the Camel’s Hump mountain in the south. The Camel’s Hump Mountains, a distinct east-west aligned range located north of the Nilgiris in the South Wayanad Hills, are recognised as an Important Bird Area (IBA) by BirdLife International, highlighting their ecological significance for avian biodiversity.

Wayanad exhibits a unique climatic profile compared to the lowland plains of Kerala. Temperatures range from 7°C during winter to 33°C in summer, with an average annual rainfall of 3,000 mm. Rainfall distribution varies significantly across the region, ranging from 2,000 mm in the drier northeast to 8,000 mm in the wetter southwest. This climatic diversity profoundly influences the region’s vegetation patterns, which include moist deciduous forests in the central and eastern areas, evergreen forests on the slopes of the Brahmagiri mountains, shola grassland complexes in the high-altitude regions, and deciduous forests along the eastern border with Karnataka.Despite a few systematic birding expeditions conducted across the forest divisions of Wayanad in the last decade, limited efforts have been made to explore the region’s unique and ecologically significant Shola Sky Islands. The forests in this region are managed by three administrative divisions under the Kerala Forest Department: the Wayanad Wildlife Sanctuary, the South Wayanad Division, and the North Wayanad Division. Biogeographically, the North and South Wayanad forest divisions are distinct, hosting some of the highest mountain ranges in the Western Ghats, with elevations ranging from 1,200 to 2,100 meters above mean sea level. These areas are characterised by their rich biodiversity, unique vegetation types, and critical ecological functions.

### 2.1. Survey Design and Field Protocol

The study was conducted from 18th to 19th January 2025 across five mountain ranges in the Western Ghats of India: Bansura, Camel’s Hump, Brahmagiri, Kurichyarmala, and Manikunnu. Surveys focused on two distinct habitat types, Shola evergreen forests and Shola grasslands above 1,200 meters above mean sea level (MSL). Thirteen base stations were established in interior forest areas to facilitate access to pre-defined transects. 50 birdwatchers were organised into 15 teams (2–3 members per team). Each team surveyed pre-designated transects (3–10 km long) along established walking paths, reflecting natural terrain rather than linear straight-line routes. Two daily observation sessions were conducted: morning (8:00 AM–12:00 PM) and afternoon (3:00 PM–6:00 PM), adjusted for local light conditions. Each transect required approximately 3 hours, ensuring standardised effort across teams.

Birds were identified through direct sightings and auditory cues using binoculars (10×50 or 8×40 magnification) and verified with regional field guides. For each observation, the species name, time, number of individuals, and behavioural remarks were recorded. Vegetation structure and land use features (e.g., forest-grassland boundaries) were noted alongside avian data. Data were logged in real-time using the eBird application, generating georeferenced checklists with latitude-longitude coordinates. Only checklists above 1,200 m MSL were retained for analysis. Teams followed standardised protocols to minimise observer bias, including cross-verification of species identification and adherence to fixed transect timings. All data were cross-checked against eBird’s global database for accuracy before analysis. Surveys adhered to non-invasive practices, avoiding disturbance to wildlife and habitats. This structured approach ensured robust data collection and integration of field observations with geospatial tools, enabling comprehensive analysis of avian diversity in Shola ecosystems.

## 3. ANALYSIS

### 3.1. Spatial Analysis of Shola Landscapes

To analyse the spatial distribution of Shola landscapes, we utilised a Geographic Information System (GIS)-based approach to compute the centroid locations and inter-centroid distances of individual landscape patches. The analysis was conducted using Geopandas, Shapely, and Matplotlib in Python, enabling efficient handling of geospatial data, projection transformations, and visualisation. The analysis began with importing a shapefile representing Shola landscapes. The dataset was loaded using Geopandas, ensuring compatibility with geospatial operations. Since the raw shapefile was in a geographic coordinate system (latitude and longitude), it was reprojected to a projected coordinate system—UTM Zone 43N (EPSG: 32643)—to facilitate accurate distance measurements. This transformation converted angular measurements into metric units, reducing distortions in spatial computations. Centroids for each Shola patch were calculated using Shapely, which derived the geometric centres of individual polygons. These centroids served as reference points for computing inter-patch distances. A pairwise Euclidean distance matrix was then constructed, where each centroid was compared against all others to determine spatial separation. Distances were measured in meters and converted to kilometres for interpretability. The resulting matrix provided quantitative insights into the spatial dispersion of Shola patches, allowing for assessments of connectivity and isolation within the landscape. To visually interpret spatial relationships, the shapefile was plotted alongside its centroids. The geometries were scaled to kilometres to maintain consistency with distance measurements. Centroids were overlaid as red markers, while inter-centroid distances were illustrated using blue dashed lines. Labels representing individual patches (e.g., mountain names) were extracted from the attribute table and displayed at their respective centroid positions. This visualisation aided in understanding the spatial configuration of Shola patches, highlighting clustering patterns or potential dispersal barriers.

### 3.2. Species Composition and Diversity Analysis

To analyse species composition across sites, we performed Non-Metric Multidimensional Scaling (NMDS) using `Python`. NMDS was chosen due to its ability to handle non-normally distributed ecological data and preserve rank-based dissimilarities, making it well-suited for species abundance data.

The dataset consisted of bird species counts across the five mountain ranges. Data preprocessing involved ensuring all values were numeric and handling missing values by replacing them with zeros using the `pandas` library. The dataset was transposed to compare sites (mountain ranges) rather than species.

#### 3.2.1. Bray-Curtis Dissimilarity

The Bray-Curtis dissimilarity index was computed using the `pdist` function from `scipy.spatial.distance`, and the matrix was converted to square form using `squareform`. This index is defined as:

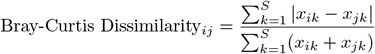

where:

- *x*_*ik*_ and *x*_*jk*_ are the abundances of species *k* at sites *i* and *j*,
- *S* is the total number of species.

#### 3.2.2. NMDS Ordination

NMDS was performed using the `MDS` function from the `sklearn.manifold` module with two dimensions and a precomputed dissimilarity matrix. The stress value, indicating the goodness of fit, was derived from the NMDS output and represents the mismatch between the distance in reduced dimensions and the original dissimilarity matrix.

#### 3.2.3. Diversity Indices

Shannon and Simpson diversity indices were computed for each mountain range to assess species richness and evenness.

*Shannon Diversity Index:* —Calculated using the entropy function from `scipy.stats`:

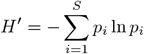

*Simpson Diversity Index:* —Calculated using:

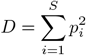

where *p*_*i*_ is the proportion of individuals belonging to species *i* and *S* is the total number of species.

#### 3.2.4. True Diversity (Hill Numbers)

To enable comparison between indices, we converted diversity values into effective numbers of species using Hill numbers.

*Shannon-based true diversity:* —Calculated using

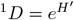

*Simpson-based true diversity:* —Calculated using:

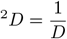

These transformations express diversity in terms of the number of equally common species that would yield the same diversity value, making them more interpretable and comparable across sites.

## 4. RESULTS

The study reported a total of 114 bird species across altitudes. The Shola forests of the South and North Wayanad regions (Camel’s Hump Mountains, Banasura Mountain, Kurichyarmala, Amaba-Vannathi Complex, and Brahmagiri) revealed a remarkable bird diversity of 114 species belonging to 44 families and 13 orders. Within the altitudinal range of 1200–1800 meters, 63 species were recorded from the Shola forests of the North Wayanad Division, and 99 species were recorded from the Shola forests of the South Wayanad Division. The diversity across mountain ranges in the Wayanad region revealed distinct bird species diversity. The Banasura Mountain range recorded 47 species, while the Brahmagiri Mountain range supported 35 species. The Camel’s Hump Mountain range, being the most biodiverse among the surveyed ranges, recorded 81 species. The Kuricharmala Mountain range had 41 species, and the Manikunnu Mountain range recorded 38 species.

The analysis of bird species distribution across different taxonomic levels revealed significant variations in species richness among families and orders. Muscicapidae exhibited the highest species count at the family level, followed by Accipitridae and Phylloscopidae, due to seasonal influence as they are migratory indicating their dominance in the observed dataset. Several families, including Campephagidae, Dicruridae, and Paridae, showed moderate species richness, while many others displayed fewer species, highlighting the uneven distribution across families. At the order level, Passeriformes overwhelmingly contained the highest number of species, far surpassing other orders, consistent with its status as the most diverse avian order globally. Other orders, such as Accipitriformes, Piciformes, and Columbiformes, had relatively lower species counts, reinforcing the dominance of passerines in avian biodiversity.

Species richness across trophic niches varies significantly among mountain ranges. Invertivores consistently dominate, with Camel’s Hump range harboring the most species (48), followed by Banasura (27), Kuricharmala (25), Manikunnu (23), and Brahmagiri (18). This pattern suggests insect prey abundance is a key driver of avian diversity. Frugivores and omnivores show moderate representation, particularly in Camel’s Hump, indicating the importance of fruit resources and dietary flexibility. Conversely, scavengers and vertivores are poorly represented across all ranges, likely due to limited carrion and vertebrate prey availability in montane ecosystems. This variation suggests that habitat heterogeneity and resource availability critically structure avian communities. Camel’s Hump, exhibiting the highest diversity, may represent a biodiversity hotspot due to its varied microhabitats, greater vegetational complexity, and intermediate anthropogenic disturbance.

A similar pattern emerges across elevational gradients, with invertivores dominating at all altitudes. Highest richness occurs at lower elevations (44 species at 1200–1300 m; 41 at 1301–1400 m), followed by gradual decline with increasing elevation. This aligns with declining arthropod abundance at higher elevations. Frugivores, omnivores, and nectarivores also decrease with elevation, reflecting shifts in floral and fruiting plant distributions. Species richness declines sharply above 1600 m, with only five invertivore species at 1700–1800 m, suggesting climatic conditions, reduced habitat complexity, and limited resources constrain diversity above this threshold. This pattern matches global montane biodiversity trends where mid-elevation zones exhibit peak richness due to optimal resource-climate balance.

The diversity indices further supported these findings (Table 1). The Camel’s Hump Mountain range exhibited the highest species richness, with 81 species, and the highest Shannon diversity index (3.2278), indicating greater species evenness and diversity. In contrast, the Banasura range had the lowest number of species (47) but maintained a relatively high Shannon diversity (3.0085), suggesting a balanced species distribution despite lower richness. The Simpson diversity index, which measures dominance, was highest for Brahmagiri (0.9313) and Camel’s Hump (0.9305), indicating lower dominance and higher evenness in these ranges.

**Table 1.**
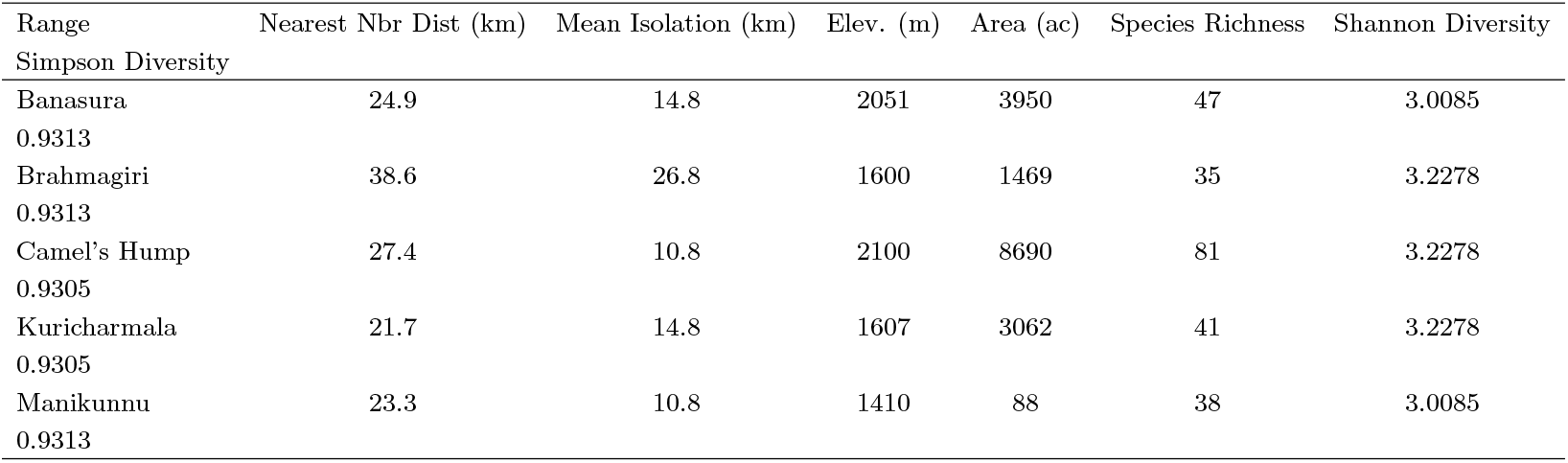
Geographic and Diversity Metrics Across Mountain Ranges in Wayanad.

The Bray-Curtis dissimilarity matrix highlighted the varying degrees of dissimilarity between the mountain ranges (Table 2). The Camel’s Hump Mountain range exhibited the highest dissimilarity values compared to other ranges, with values ranging from 0.6928 to 0.7934. In contrast, the Banasura and Brahmagiri ranges showed relatively lower dissimilarity (0.4834), suggesting more similar bird communities between these two ranges. The Non-Metric Multidimensional Scaling (NMDS) ordination, with a stress value of 0.1461, provided a robust visualization of the dissimilarity patterns among the mountain ranges (Figure 6). The stress value indicated a good fit, as values below 0.2 are generally considered acceptable in ecological studies. The NMDS plot revealed a clear separation of the Camel’s Hump Mountain range from the other ranges, suggesting a distinct bird community structure in this region. In contrast, the Banasura and Brahmagiri ranges clustered more closely, indicating greater similarity in their bird species composition.

**Table 2.** Pairwise Isolation Index Among Mountain Ranges in Wayanad. Values represent scaled distances between range centroids (unitless).

| Mountain Range | Banasura | Brahmagiri | Camel's Hump | Kuricharmala | Manikunnu |
| --- | --- | --- | --- | --- | --- |
| Banasura | 0.0000 | 0.4834 | 0.6928 | 0.5204 | 0.5574 |
| Brahmagiri | 0.4834 | 0.0000 | 0.7934 | 0.5689 | 0.5182 |
| Camel's Hump | 0.6928 | 0.7934 | 0.0000 | 0.7583 | 0.7664 |
| Kuricharmala | 0.5204 | 0.5689 | 0.7583 | 0.0000 | 0.5803 |
| Manikunnu | 0.5574 | 0.5182 | 0.7664 | 0.5803 | 0.0000 |

**Table 3.**
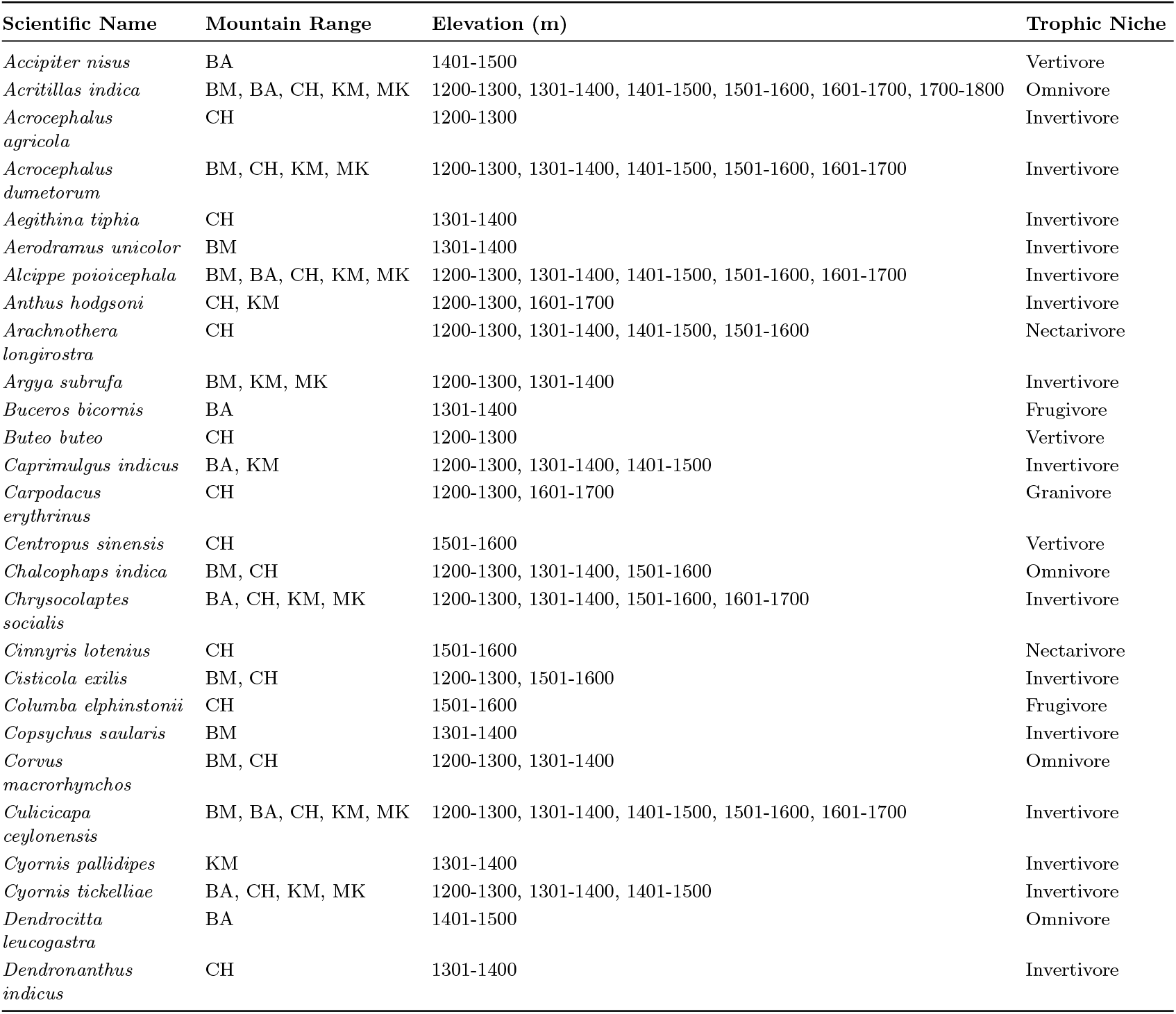
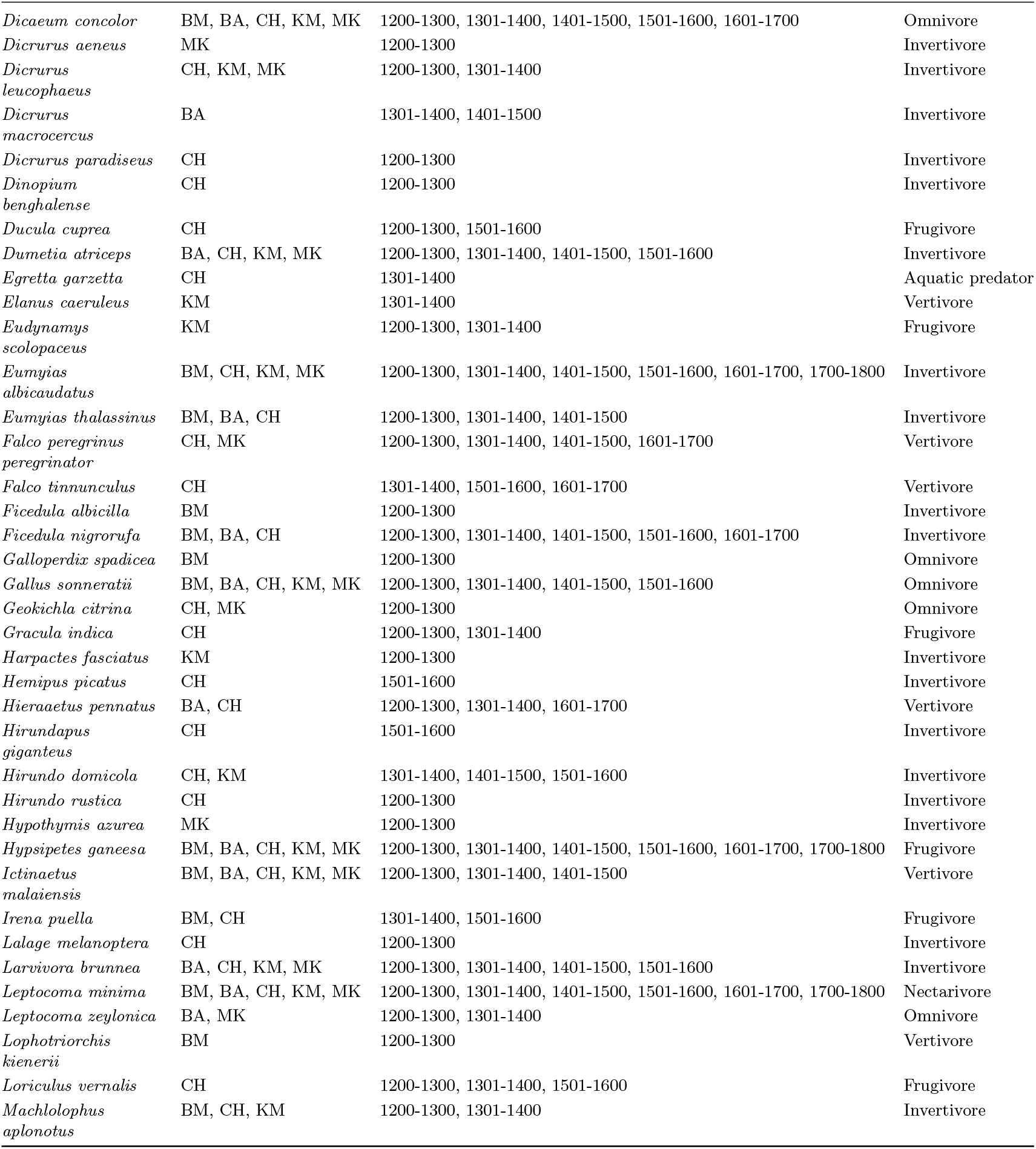
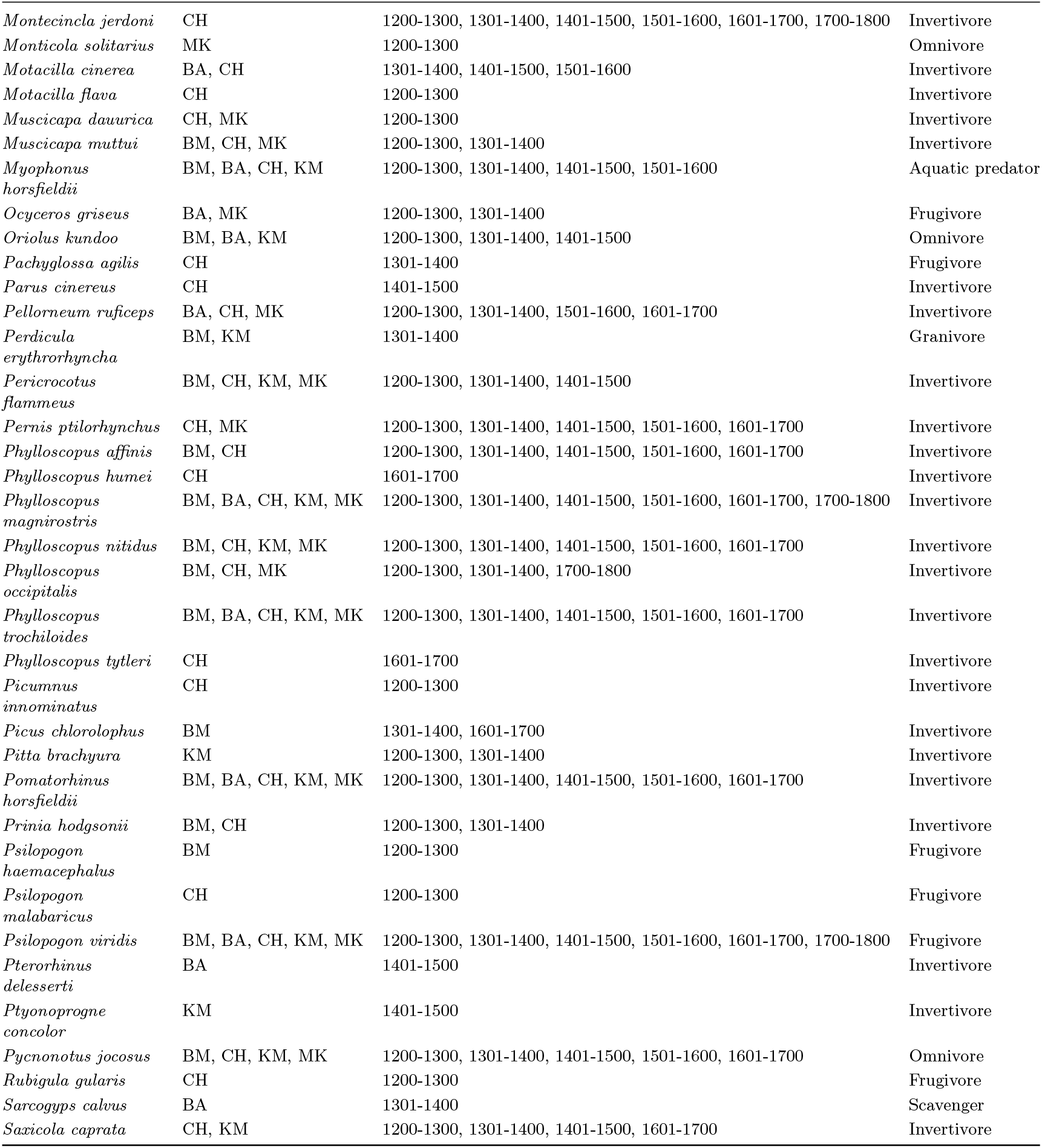
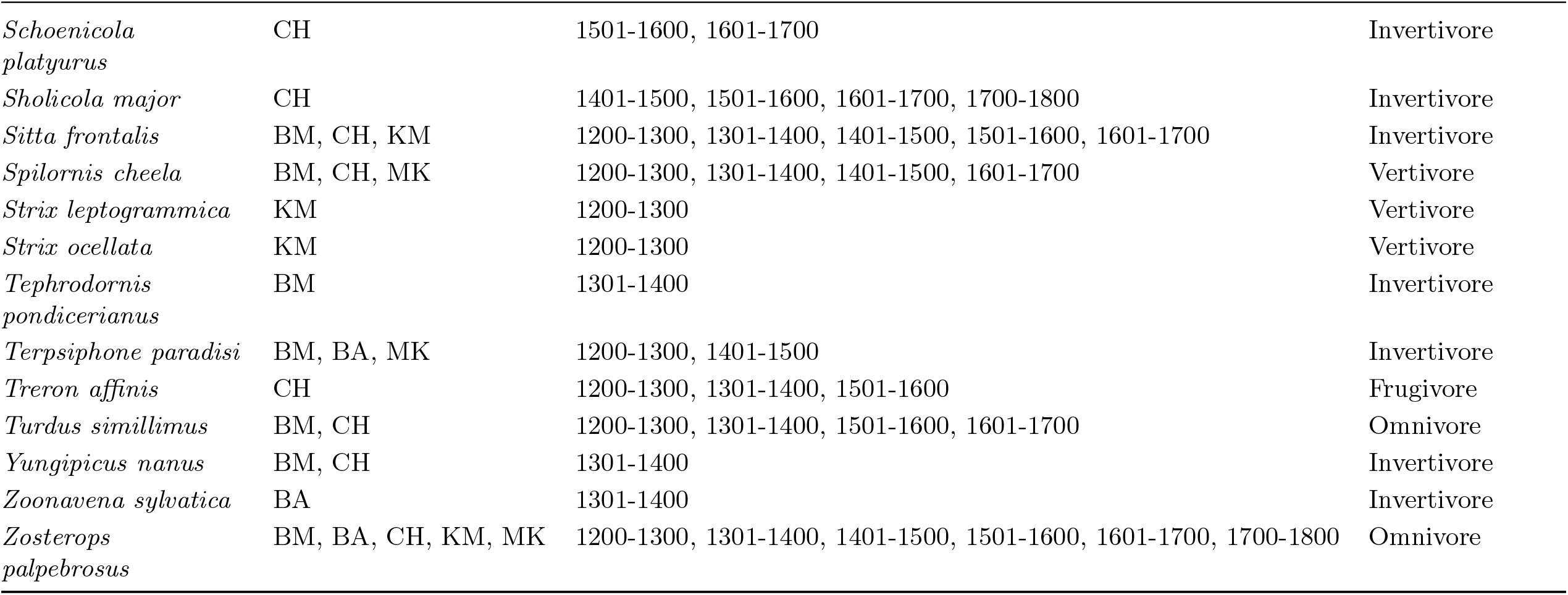
Bird Species Data from Wayanad Mountain Ranges (BM=Banasura, BA=Brahmagiri, CH=Camel’s Hump, KM=Kuricharmala, MK=Manikunnu)

**Figure 1.**
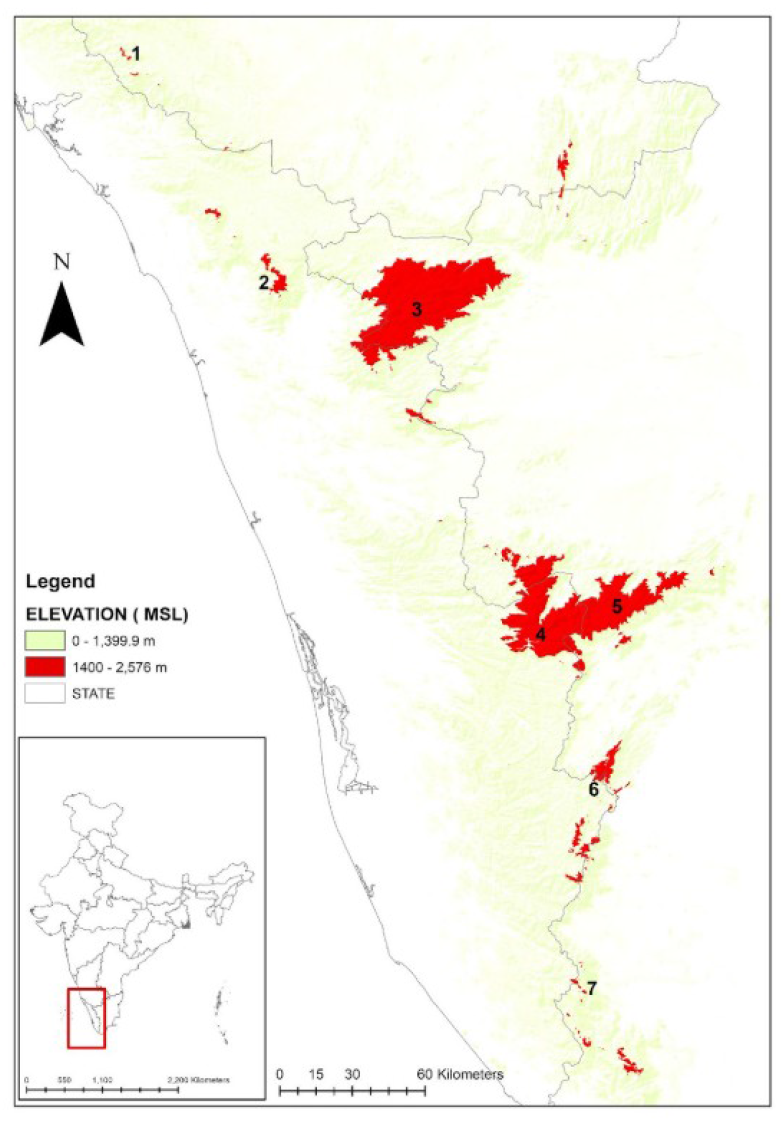
The Shola Sky Islands of the Western Ghats consist of several significant mountain-tops (1 to 7). The northern sky islands include Brahmagiri (1) and Wayanad (2). The Baba Budan Hills, Kudremukh, and Pushpagiri are other mountains from the Northern Sky islands that are not plotted on the map. The highest elevation plateaus form the central sky islands, comprised of the Nilgiri mountains (3) and across the deep Palghat Gap, the Anamalai Hills (4), and Palani Hills (5), which form a single high-elevation plateau. C: The southern sky islands comprise the Meghamalai Hills (6) and the Agasthyamalai or Ashambu Hills (7) mountain range.

**Figure 2.**
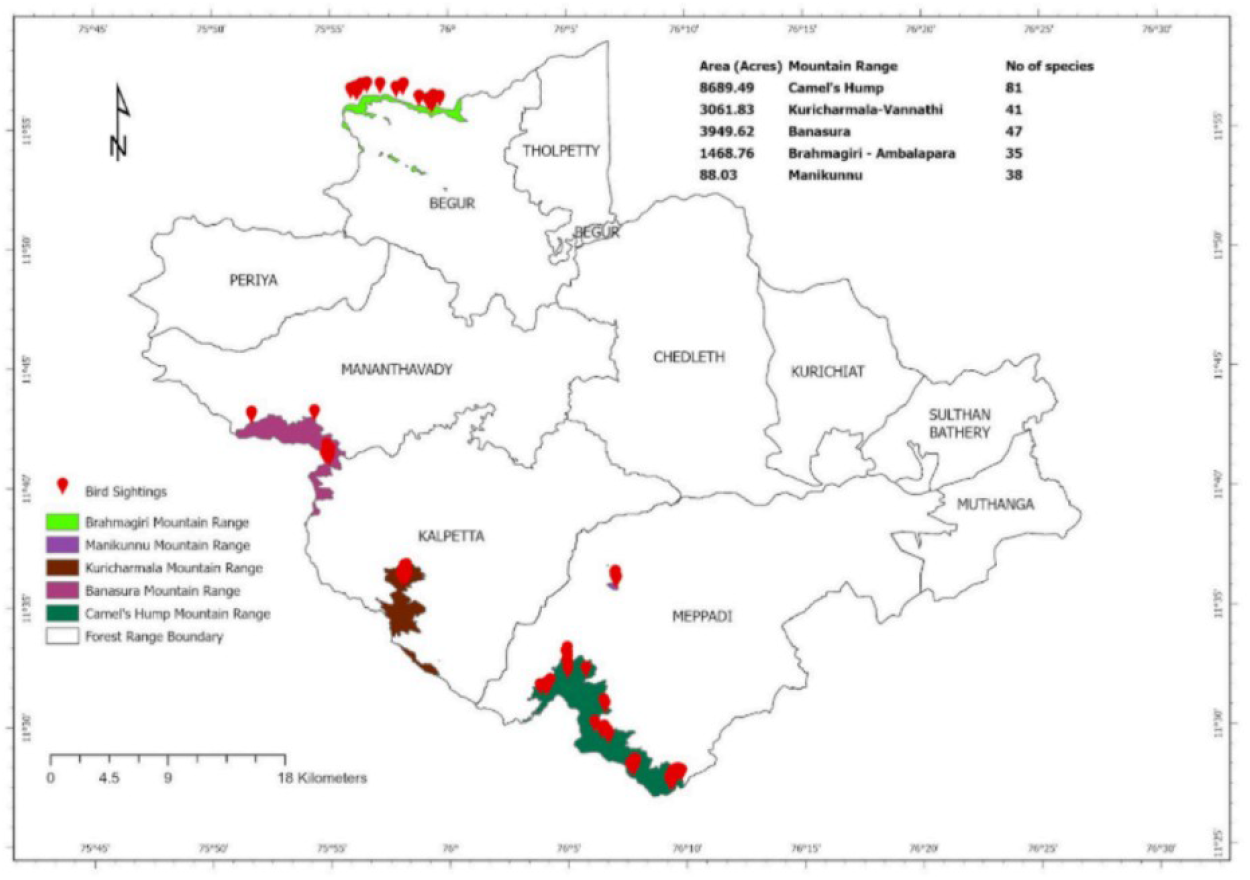
Map showing the distribution of eBird checklists submitted in Wayanad district, Kerala, India, from January 2023 to December 2023, illustrating spatial patterns of bird observation records.

**Figure 3.**
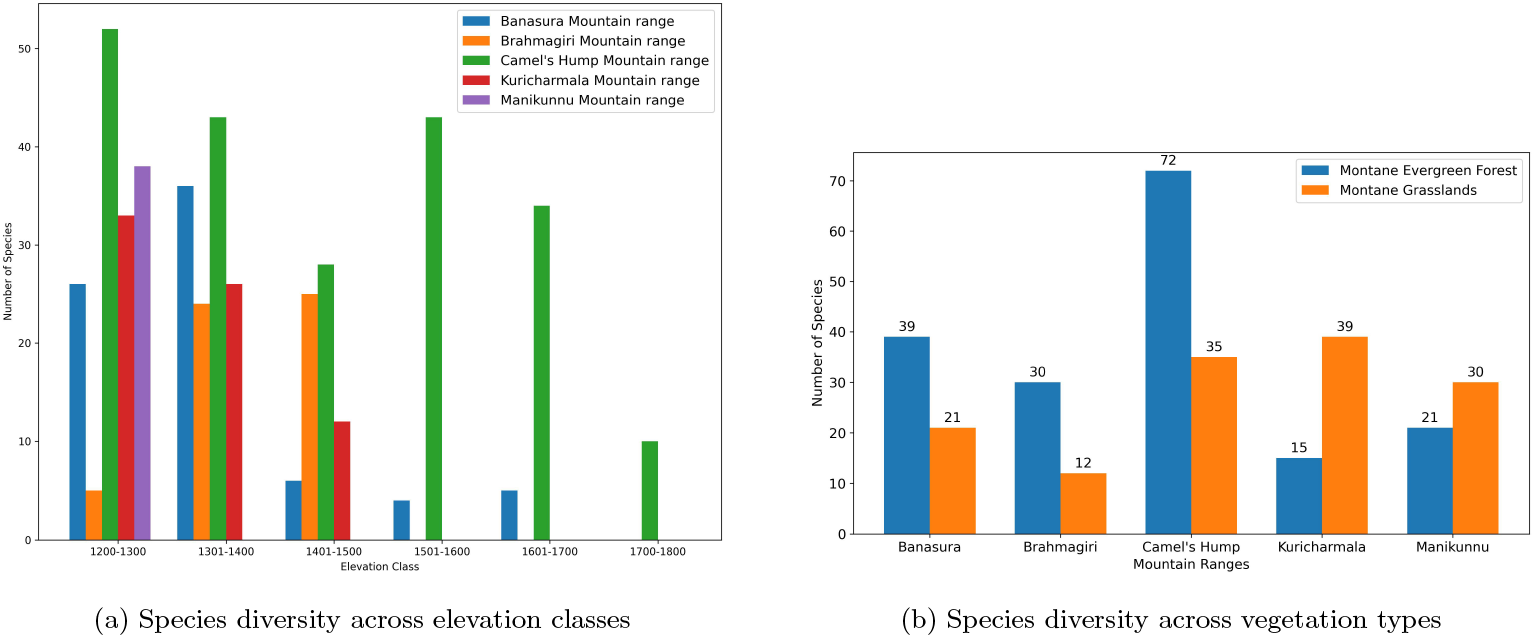
(a) Bird species distribution across elevation classes. (b) Species counts in two vegetation types across the mountain ranges.

**Figure 4.**
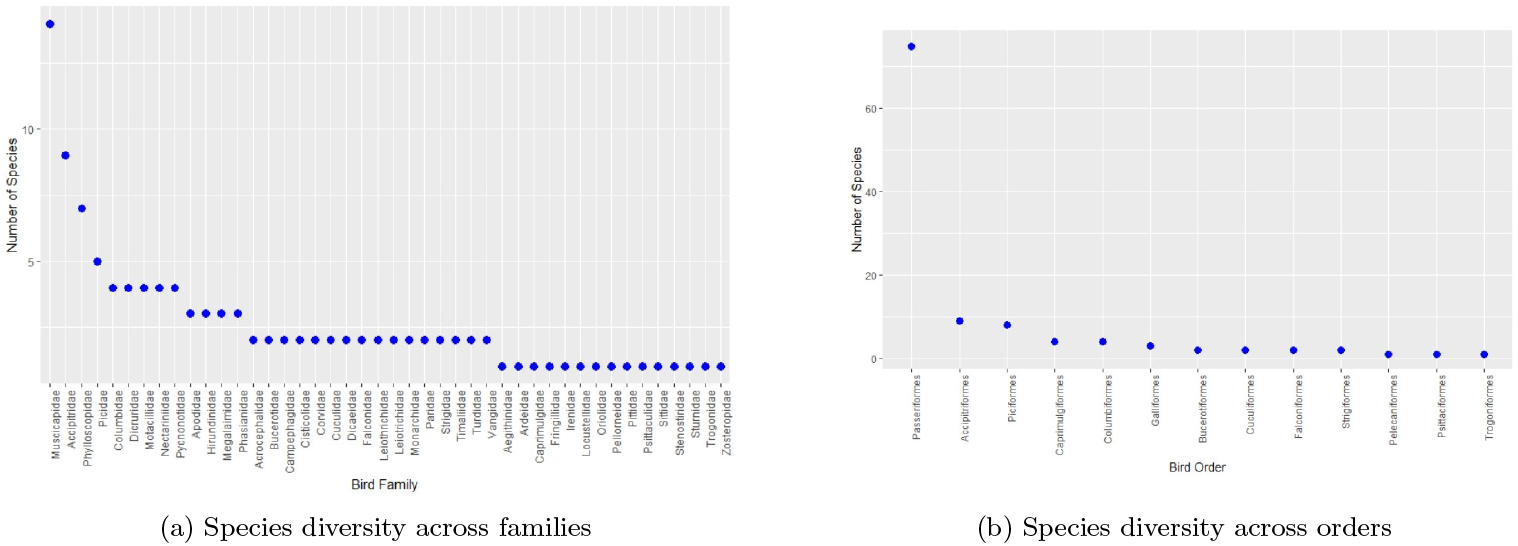
(a) Bar graph showing number of bird species across families, with Muscicapidae having the highest count followed by Accipitridae and Phylloscopidae. (b) Bar graph showing species distribution across orders, with Passeriformes as the most species-rich order.

**Figure 5.**
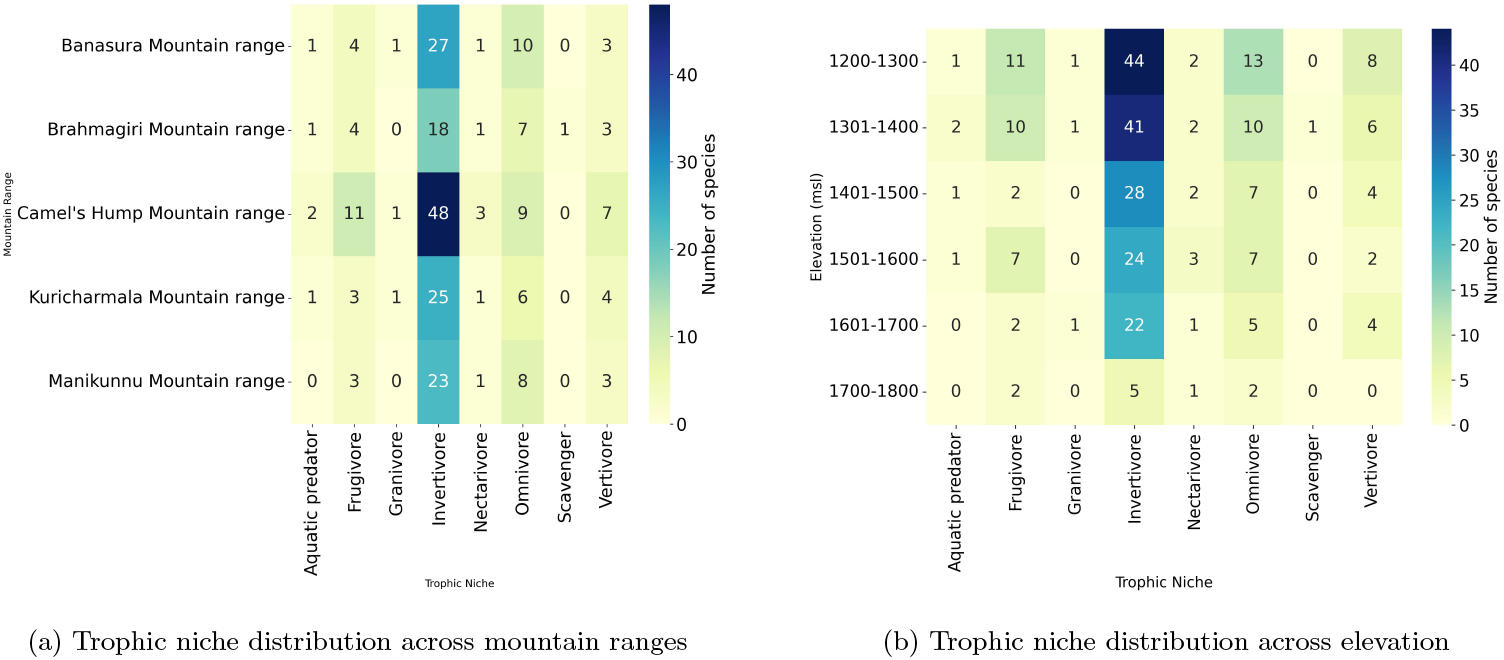
(a) Trophic niche distribution across five Wayanad mountain ranges. Invertivores dominate all ranges, particularly Camel’s Hump. (b) Heatmap showing species distribution across trophic niches and elevation bands (1200–1800 m). Invertivores dominate all elevations with highest richness at 1200–1500 m.

**Figure 6.**
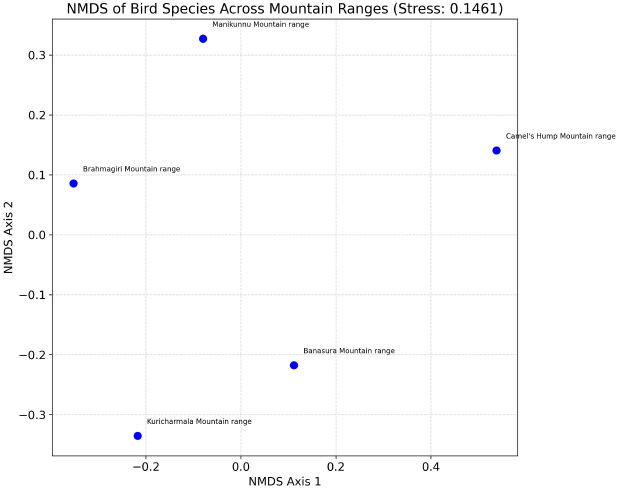
NMDS ordination showing bird species composition across five mountain ranges in Wayanad (Bray-Curtis dissimilarity; Stress = 0.1461). Spatial separation indicates compositional differences among ranges.

**Figure 7.**
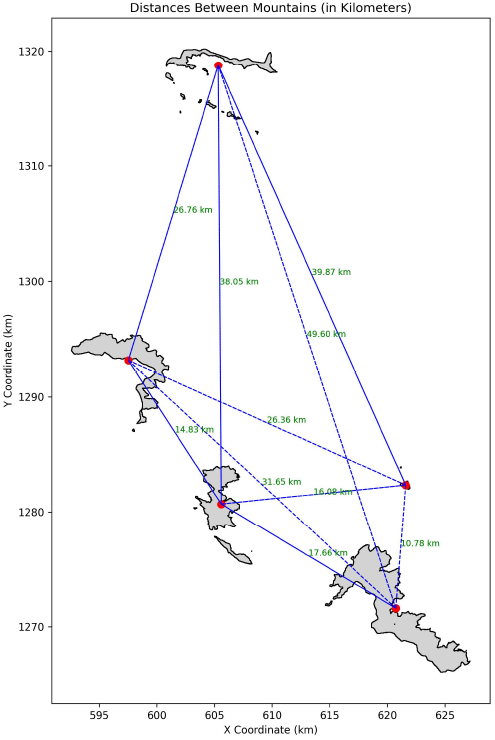
Distance between key mountain ranges in the study area.

This study of five Shola sky islands in the Wayanad demonstrates that biodiversity patterns align strongly with core tenets of island biogeography theory. Consistent with the species-area effect, larger islands like Camel’s Hump (8,689.49 acres) hosted significantly higher species richness (81 species) and diversity indices (Shannon=25.22, Simpson=14.39) compared to smaller islands like Manikunnu (88.03 acres; 38 species) or Brahmagiri (1,468.76 acres; 35 species). Furthermore, the distance effect was evident: Brahmagiri, the most isolated patch (38.57 km to nearest neighbour), recorded the lowest richness (35 species), while islands with closer proximity, such as Kuricharmala and Banasura (14.83 km apart), supported higher richness (41 and 47 species respectively). The data revealed key interactions; Camel’s Hump’s large area buffered its moderate isolation (mean 27.42 km), sustaining high diversity, while Manikunnu’s proximity to Camel’s Hump (10.78 km) mitigated its small size, resulting in moderate diversity (Shan- non =3.02). Despite subtle elevation variations, shared Shola habitat minimised confounding effects, emphasising area and isolation as the dominant drivers. These findings confirm the applicability of MacArthur and Wilson’s theory to montane sky islands, highlighting larger islands as biodiversity reservoirs and proximity as crucial for dispersal. Consequently, conservation must prioritize protecting large islands (e.g., Camel’s Hump) and enhancing connectivity between proximate patches (e.g., Banasura-Kuricharmala), while implementing targeted strategies for small, isolated islands like Brahmagiri, acknowledging context-specific modifiers like climatic stability and topographic barriers within these terrestrial “island” systems.

## APPENDIX

