## Supplementary figures and images for "Wayanad Sky Islands Survey 2025"

### elevation map.jpg

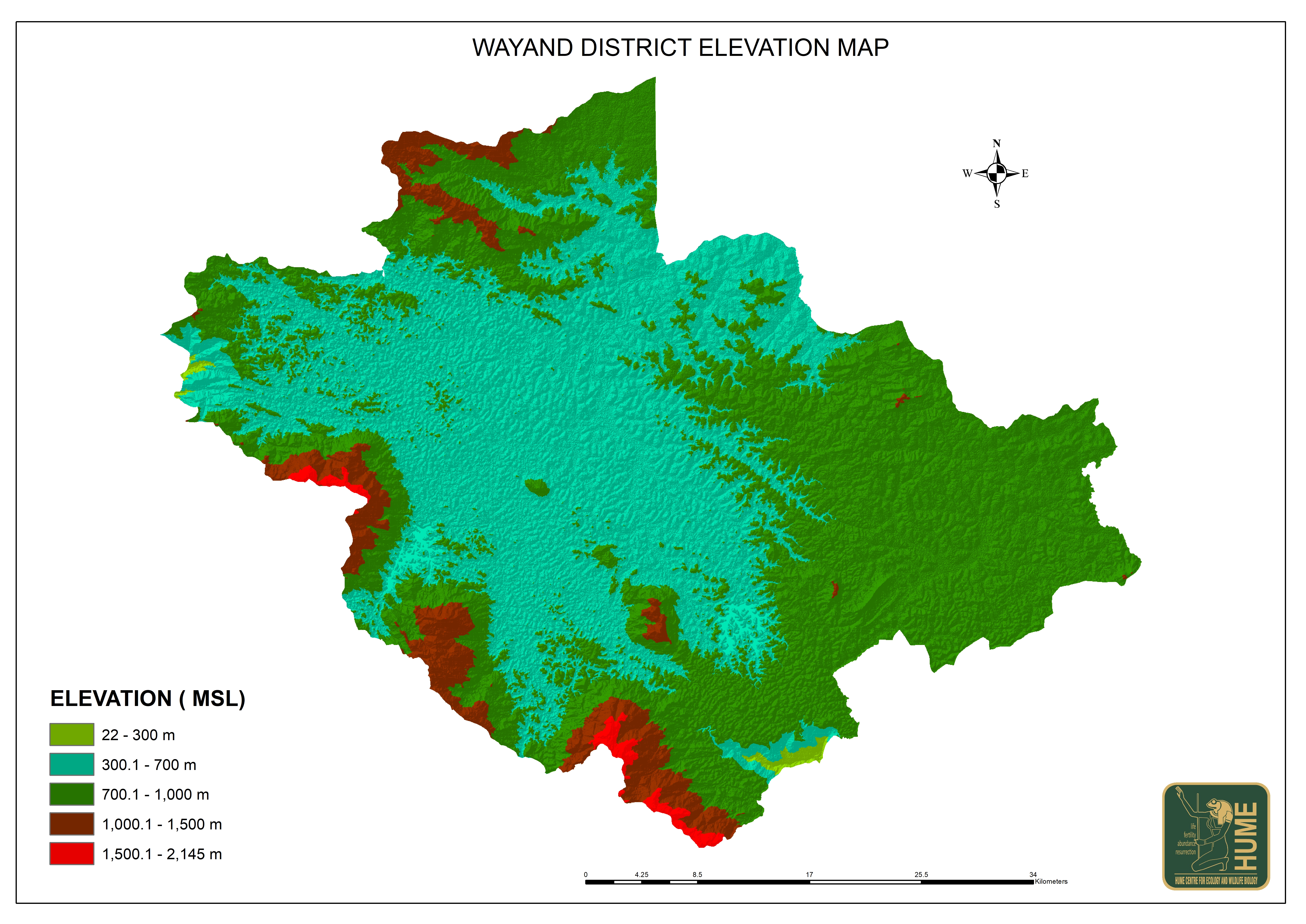

### Mountain_range_network.jpeg

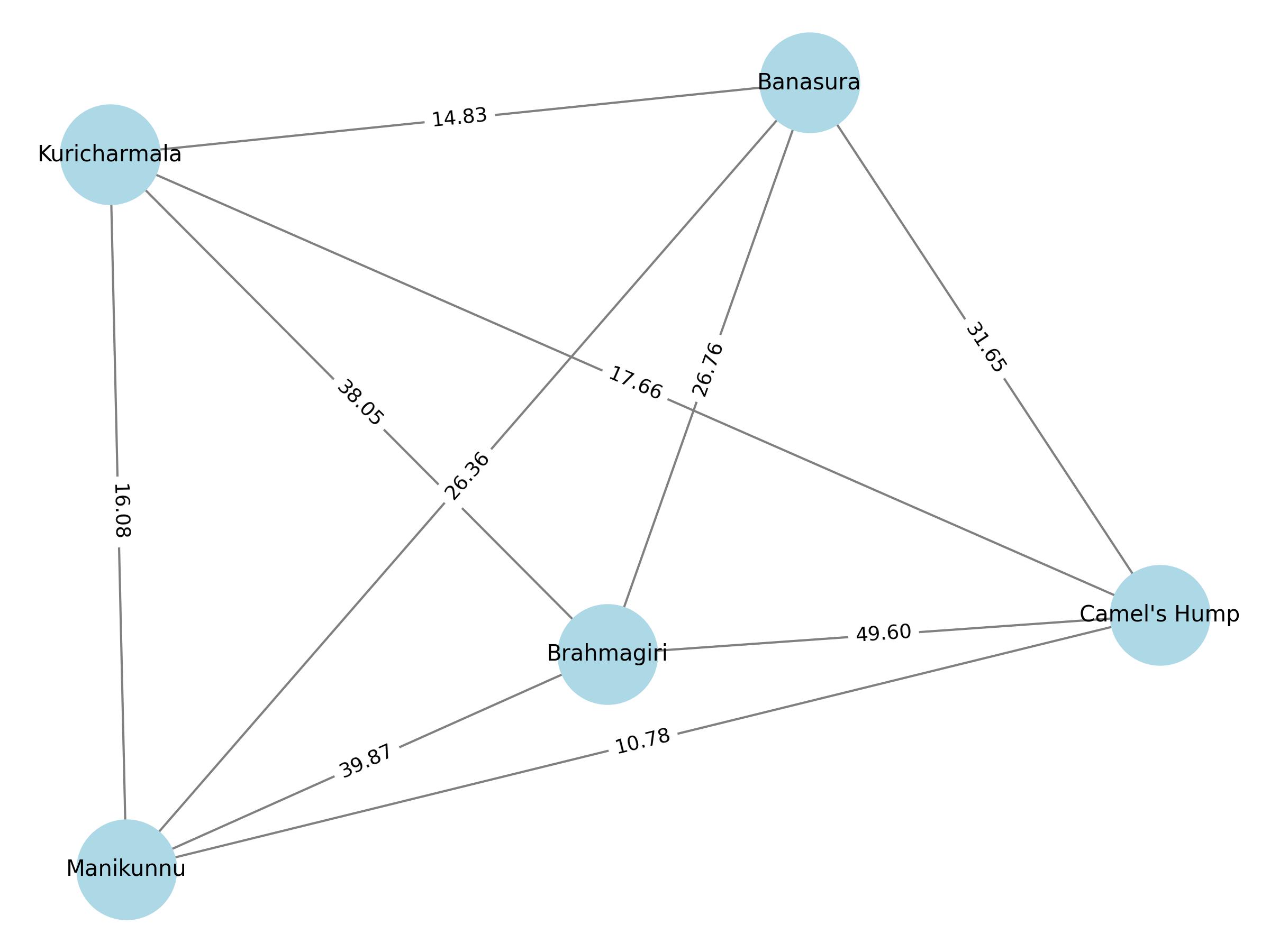
